# Auditory cortex supports verbal working memory capacity

**DOI:** 10.1101/2020.08.05.237727

**Authors:** Gavin M. Bidelman, Jane A. Brown, Pouya Bashivan

## Abstract

Working memory (WM) is a fundamental construct of human cognition. The neural basis of auditory WM is thought to reflect a distributed brain network consisting of canonical memory and central executive brain regions including frontal lobe, prefrontal areas, and hippocampus. Yet, the role of auditory (sensory) cortex in supporting active memory representations remains controversial. Here, we recorded neuroelectric activity via EEG as listeners actively performed an auditory version of the Sternberg memory task. Memory load was taxed by parametrically manipulating the number of auditory tokens (letter sounds) held in memory. Source analysis of scalp potentials showed that sustained neural activity maintained in auditory cortex (AC) prior to memory retrieval closely scaled with behavioral performance. Brain-behavior correlations revealed lateralized modulations in left (but not right) AC predicted individual differences in auditory WM capacity. Our findings confirm a prominent role of auditory cortex, traditionally viewed as a sensory-perceptual processor, in actively maintaining memory traces and dictating individual differences in behavioral WM limits.

## Introduction

Working memory (WM) is the mental process that temporarily preserves and manipulates information for later deployment to our perceptual-cognitive systems. WM operations consist of both memory retrieval and active manipulations of sensory-cognitive representations. Human WM is, however, a limited capacity system (Cowan 2001), capable of buffering ~7±2 items in memory store at any one time (Miller 1956). Given its ubiquity to perceptual-cognitive processing, defining the neural mechanisms of WM is important to understand the brain basis of this core cognitive function.

Neuroimaging studies have identified correlates of WM in canonical memory and central executive brain regions including parietal lobe, (pre)frontal areas, and hippocampus (Bashivan et al. 2014b; Karlsgodt et al. 2005; Kumar et al. 2016). Cross-species studies corroborate human work by implicating higher cognitive association areas outside sensory cortices (Constantinidis et al. 2004). Moreover, different stages of WM (encoding, maintenance, and retrieval) recruit different neural circuitry (Bashivan et al. 2014a; Karlsgodt et al. 2005). In the context of visual WM, we have recently shown the directional flow of information between sensory and frontal brain areas reverses when encoding vs. maintaining items in WM (*encoding*: occipital sensory→frontal; *maintenance*: frontal→occipital sensory), revealing feedforward and feedback modes in the same underlying brain network (Bashivan et al. 2014a). While lower-(sensory) and higher-level (cognitive) brain regions interact during the time course of WM processing, an outstanding question to address is the degree to which sensory cortex itself accounts for differences in WM capacity.

A persistent and dominant view is that regions beyond auditory cortex (AC) drive WM in human and non-human primates (Grady et al. 2008; Huang et al. 2016; Kumar et al. 2016; Lefebvre et al. 2013). Thus, while AC is responsible for precise stimulus encoding, it may not itself participate in the active maintenance of information in a memory buffer. Retrieval and subsequent manipulation of mental objects would be orchestrated via controlled coupling between sensory cortex and frontal and/or hippocampal areas (Boran et al. 2019). Critically, such models assume that regions beyond Heschl’s gyrus maintain information, although perhaps in a different representational form (Yue et al. 2019).

On the contrary, emerging evidence suggests a substantial portion of auditory WM might indeed be supported by more automatic, lower-level processing *within* the auditory system without the need for (or minimal reliance on) higher-level brain structures (Linke et al. 2011). Evidence across species suggests activity in superior temporal gyrus (STG) (Acheson et al. 2011; Grimault et al. 2014) predicts listeners’ WM capacity. However, involvement of STG/STS in verbal WM, particularly posterior regions (e.g., Acheson et al. 2011), might be expected given the role of these “dorsal stream” areas in phonological production, articulatory planning, and auditory-motor transformation (Hickok et al. 2007). Growing evidence implicates sensory AC itself in WM processing (Grady et al. 2008). In non-human mammals, AC neurons show sustained activity that is sensitive to WM demands (Sakurai 1994). Such findings suggest that auditory WM, a function traditionally viewed through a “cognitive lens,” might be driven not by higher-order neocortex, *per se*, but nascent memory representations in auditory areas (e.g., Kumar et al. 2016).

To investigate the role of AC in auditory WM, we recorded EEG as listeners performed an serial auditory WM task. We varied WM demand (i.e., cognitive load) by manipulating the number of tokens (letter sounds) held in memory. Using source analysis, we examined the underlying brain regions modulated by memory load and, more critically, whether neural activity prior to memory retrieval could predict listeners’ subsequent behavioral performance. Our findings reveal early stages of auditory cortical processing in left hemisphere reflect a direct correlate of the auditory WM trace and robustly predict individual differences in behavioral capacity limits.

## Methods

### Participants

The sample included *n*=15 young adults (28±3 yrs; 8 female) recruited from the University of Memphis. All had normal hearing (thresholds < 25 dB HL), reported no history of neuropsychiatric illness, and were right-handed (Edinburgh laterality index > 95%). One participant’s data was excluded due to excessive noisy EEG. Each gave written informed consent in compliance with a protocol approved by the University of Memphis IRB.

### Stimuli and task

EEGs were recorded during a version of the Sternberg memory task that parametrically varied memory load while temporally separates *encoding, maintenance*, and *recall* stages of WM processing (for details, see Bashivan et al. 2014b). On each trial, listeners heard a random series of alphanumeric characters from a subset of 15 English letters (**Fig. 1a**). Tokens were 300 ms (SOA = 1000 ms) presented at 76 dB SPL over ER-3A headphones (Etymotic Research). For each trial, we varied the length (size) of the “Memory set” 2, 4, 6, or 8 characters (random draw). A 3000 ms delay (“Retention”) followed the offset of the final stimulus in which listeners retained the set in memory. Following retention, a single “Probe” character was played and participants indicated if it was among the prior memory set. Feedback was provided after 300 ms. The next trial commenced after 3.4 sec. On half the trials, the probe occurred in the set; on the other half it did not. Participants received 20 practice trials for familiarization and completed a total of 60 experimental trials per set size. The task was coded in MATLAB using the Psychophysics Toolbox (Brainard 1997). We logged both accuracy (% correct) and median reaction times (RTs) for probe recall. We computed listeners’ WM capacity (per set size) using the *K* = *S*(*H-F*), where *S* is the number of items in the memory array, *H* is the hit rate, and *F* is the false alarm rate (Bashivan et al. 2014b; Cowan 2001).

### EEG recordings

EEGs were recorded from 64 channels at standard 10-10 locations. Continuous data were digitized at 500 Hz with a DC-250 Hz passband (SynAmps RT amplifiers; Compumedics Neuroscan). Electrodes on the outer canthi and the superior and inferior orbit monitored ocular movements. Impedances were < 5 kΩ. Response were common average referenced.

Preprocessing was conducted in BESA^®^ Research (v7) (BESA GmbH) and FieldTrip (Oostenveld et al. 2011). Blinks were nullified using principal component analysis (PCA). Cleaned EEGs were filtered (0.01-20 Hz; 20^th^ order elliptical), epoched −6000 to +4000 ms (where *t*=0 is probe onset), and averaged per set size and listener. This window encapsulated transient ERPs to the last memory tokens in the encoding period, sustained activity within the maintenance period of interest, and subsequent retrieval/response-related ERPs following the probe (see **Fig. 2**). We baselined traces between −4400 and −3700—zeroing EEGs just prior to the final memory set token—to highlight neural activity during maintenance relative to the immediately preceding encoding period.

### ERP sensor and source analysis

From channel-level waveforms, we measured ERP amplitudes within a cluster of central electrodes (C1/2, Cz, CP1/2, CPz) where load-dependent changes were prominent in initial visualizations of scalp topographies (data not shown). We then measured the mean ERP amplitude in the entire maintenance window (−3000-0 ms) within this electrode cluster (**Fig. 2,** *inset*).

We used Classical Low Resolution Electromagnetic Tomography Analysis Recursively Applied (CLARA) [BESA^®^ (v7)] to estimate the neuronal current density underlying WM. CLARA renders focal source images by iteratively reducing the source space during repeated estimations. On each iteration (x2), a spatially smoothed LORETA image was recomputed.

Voxels below 1% max amplitude were removed. Two iterations were used with a voxel size of 7 mm in Talairach space (regularization = 0.01% singular value decomposition). Group-level statistical (*t*-stat) maps were computed on full brain volumes using the ‘ft_sourcestatistics’ function in FieldTrip (Oostenveld et al. 2011) (*P*<0.05 mask). From individual CLARA images, we extracted source amplitudes at the midpoint of the maintenance period (i.e., latency of −1500 ms) within the spatial centroids of significant clusters within bilateral superior temporal gyrus (i.e., AC) (see **Fig. 3a**).

### Statistical analysis

We used mixed-model ANOVAs in R (R Core team, 2018; *lme4* package) with a fixed effect of set size (subjects = random effect). We used Tukey-Kramer corrections for multiple comparisons. To assess relations between region-specific neural activity and behavior, we regressed the change in AC source activation (i.e., set size 8 – set size 2) with individuals’ maximum WM capacity for the highest load condition (*K_load 8_*) (e.g., Grimault et al. 2014). Left and right hemispheres were analyzed separately. Robust regression (bisquare weighting) was performed using the ‘fitlm’ function in MATLAB^®^ 2019b (The MathWorks, Inc.).

## Results

### Behavioral data

Recall accuracy expectedly declined [*F*_3,39_=23.60, *p*<0.0001] and RTs slowed [*F*_3,39_=11.31, *P*<0.0001] for increasing memory load (**Fig. 1b,c**). Similarly, *K*, an unbiased measure of WM capacity, increased with set size [*F*_3,39_=48.95, *P*<0.0001] but with more dramatic improvements between 2 and 4 items (*t*_39_=−5.62, *P*<0.0001) and capacity plateauing between 6 to 8 items (*t*_39_=−2.89, *P*=0.0303) (**Fig. 1d**). For 8 stimulus items, *K* capacity was 4.72 ± 1.07, indicating ~4-5 items could be adequately maintained in auditory WM (Vogel et al. 2004). This is consistent with known limits to WM capacity (cf. 7±2; Miller 1956) observed in both the visual and auditory modalities (Bashivan et al. 2014b; Cowan 2001; Grimault et al. 2014; Lefebvre et al. 2013; Vogel and Machizawa 2004).

**Fig. 1:**
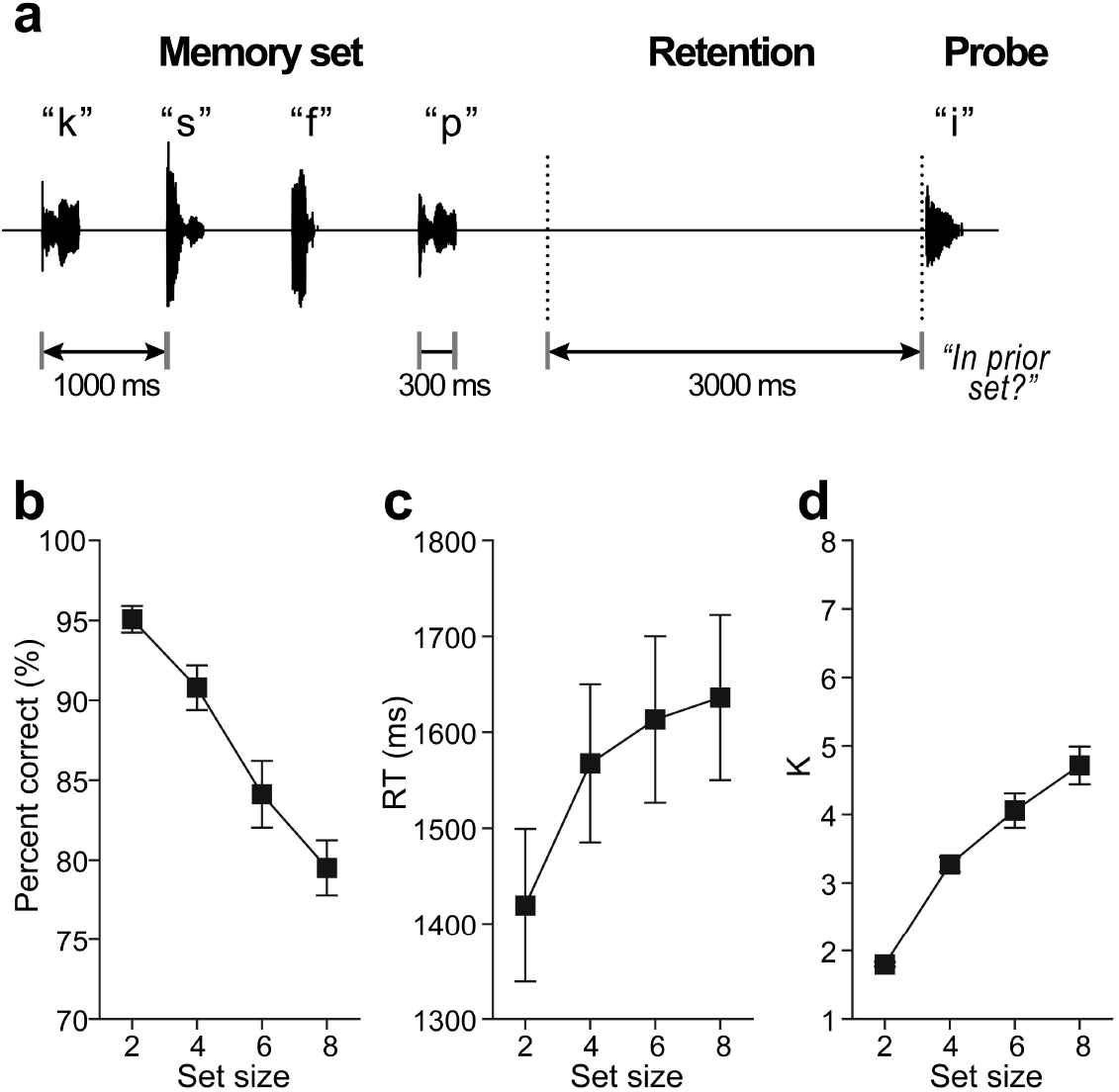
Auditory WM stimulus paradigm and behavioral data. (**a**) Listeners heard between 2 and 8 characters (“Memory set”) presented auditorily. Following a retention period, they indicated whether a “Probe” occurred in the prior memory set. Shown here is a “no match” trial. (**b-c**) Behavioral accuracy for probe recall decreases and response times increase with additional memory load. (**d**) Working memory capacity (*K*) increases for small set sizes but saturates >4 items, above listeners’ WM capacity limits. errorbars =± 1 s.e.m.

### EEG data

The time course of scalp ERPs (**Fig. 2**) tagged distinct phases of the task including evoked peaks reflecting sensory responses to the final tokens of the stimulus array and probe. Moreover, with our baseline definition, transient activity at the end of the encoding period (i.e., ERP positivity, −4000 ms) before maintenance was similar between set sizes [*F*_3,39_=1.64, *P*=0.19]. Yet, strong load-dependent changes in neural activity were observed during maintenance [*F*_3,39_=4.98, *P*=0.0051]. Scalp ERPs were larger for lower (2/4) vs. higher (6/8) memory loads [sets size 2/4 vs. 6/8: *t*_39_=3.84, *P*=0.0004] (**Fig. 2, *inset***), resembling a fatigue of neural activity in more demanding conditions.

**Fig. 2:**
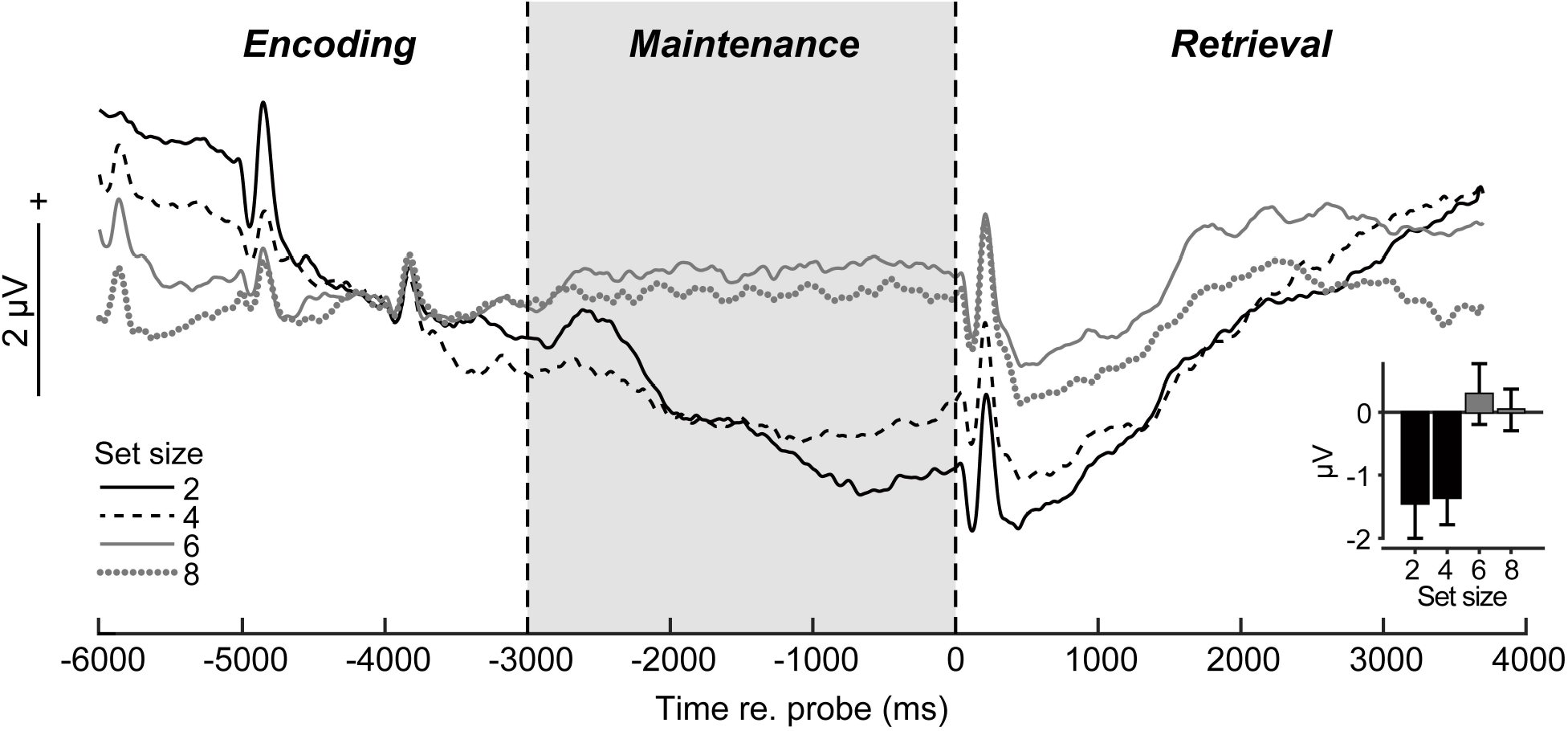
Scalp ERPs reveal load-dependent modulations in sustained neural activity during WM maintenance. **(a)** ERP time courses at central scalp locations (mean of electrodes C1/2, Cz, CP1/2, CPz; baseline = [-4400 to −3700 ms]). Transient peaks during “encoding” reflect auditory responses to final stimulus tokens in the memory set. Sustained activity is modulated in the 3 sec maintenance interval during memory retention (highlighted segment) and is stronger for low (2/4) vs. high (6/8) load (inset). errorbars =± 1 s.e.m.

Source ERP analysis uncovered activations distinguishing WM load in foci located in AC, with coverage in primary and posterior auditory cortices (BA 41/42 and BA 22) (**Fig. 3a**). AC amplitudes varied with hemisphere [*F*_1,91_=4.76, *P*=0.0317] and set size [*F*_3,91_=4.78, *P*=0.0038] with no interaction [*F*_3,91_=0.65, *P*=0.58]. Responses were stronger in left compared to right hemisphere and AC differentiated lower (2/4) vs. higher (6/8) load conditions (**Fig. 3b,c**).

Lastly, brain-behavioral regressions revealed listeners’ maximum WM capacity (i.e., *K_load 8_*) was predicted by load-dependent changes in source responses for left [*R^2^*= 0.36, *P*=0.0401] but not right AC [*R^2^*= 0.45, *P*=0.37] (**Fig. 3d,e**).

**Fig. 3:**
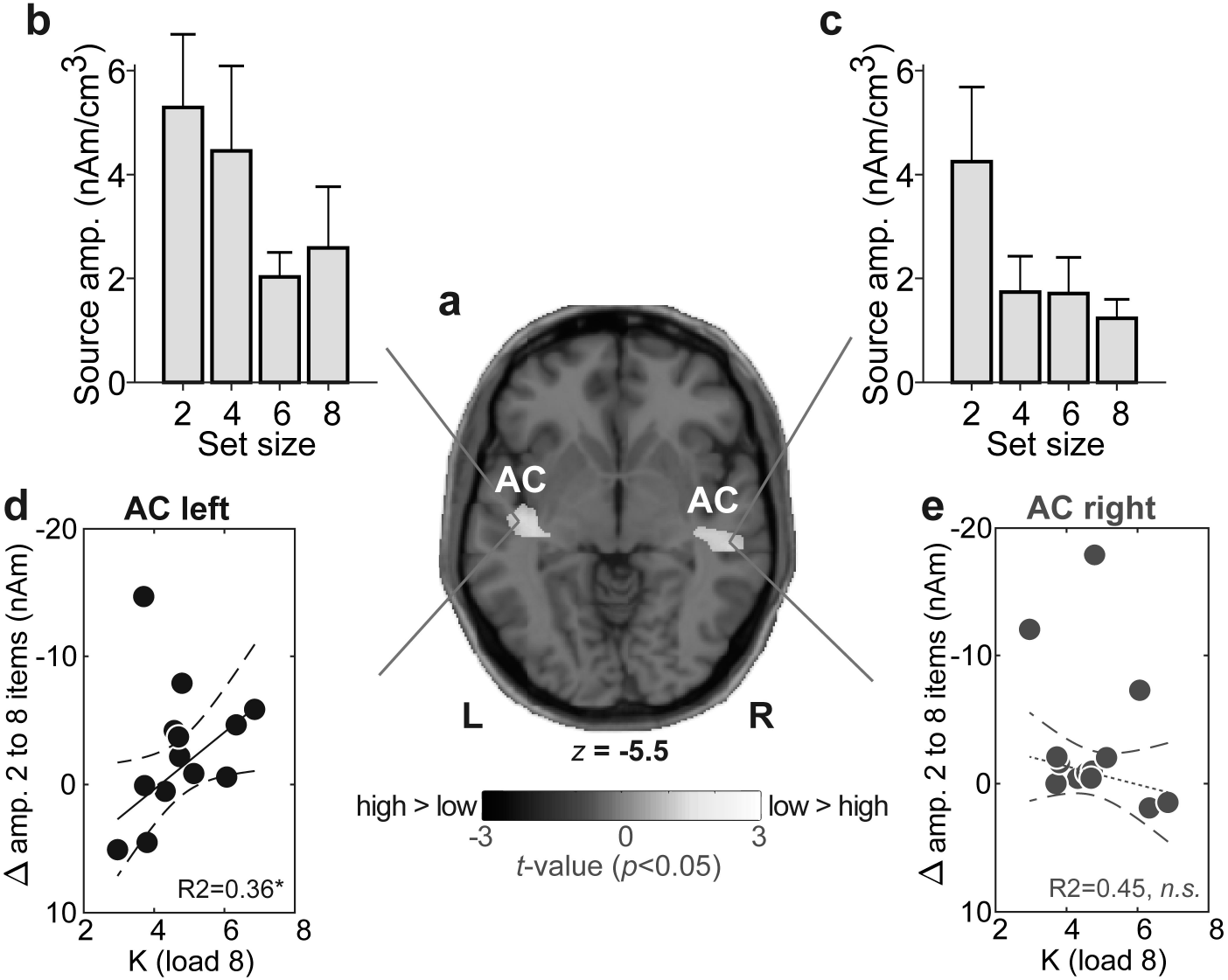
Sustained neural activity maintained in AC predicts behavioral auditory WM capacity. **(a)** T-stat map contrasting low (2/4) vs. high (6/8) CLARA source activation maps (*P*<0.05 masked, uncorrected). Functional data are overlaid on the MNI brain template. WM load is distinguished in bilateral auditory AC [MNI coordinates (*x,y,z*; in mm): AC_*left*_=(−42.5, −18.5, −5.5); *AC_right_*=(50.5, −26.5, −5.5)] **(b-c)** AC amplitudes vary with set size but load-related changes in left (but not right) AC mirror the pattern observed in scalp EEG (cf. Fig. 2a) (**d-e**) Maximum behavioral WM capacity (*K_load 8_*) is predicted by AC activity in left hemisphere; individuals with larger changes in source amplitudes with set size have larger WM capacity. No brain-behavior relation is observed in right hemisphere. Dashed lines=95% CI; solid lines, significant correlation; dotted lines, n.s. correlation. \**P*<0.05. errorbars =± 1 s.e.m.; AC, auditory cortex.

## Discussion

Using EEG during active WM tasks, we show early stages of auditory cortical processing in AC reflect a robust neural correlate of the auditory WM trace. This neural index of WM performance is dominant in left hemisphere where the degree of modulation in AC predicts individuals’ behavioral WM capacity (i.e., larger ERP changes associated with larger memory store).

Source localized activity showed load-related modulations at both the sensor and source level with larger sustained responses observed for lower- (easier) compared to higher-load (harder) set size conditions. Previous fMRI work linking sustained delay period activity to WM performance is equivocal. Some reports show enhancements (Kumar et al. 2016) and others suppression (Linke et al. 2011) of cortical responses with memory load. ERP studies are similarly ambiguous finding decreases and increases (Grimault et al. 2014; Huang et al. 2016; Lefebvre et al. 2013; Vogel and Machizawa 2004) in late (> 400 ms) slow wave potentials with load level. Reconciling these findings, direct recordings in animals show *both* suppression and enhancement effects in AC neurons dependent on the mnemonic context of sounds (Scott et al. 2014).

Sustained firing patterns might be not be related to WM *per se*, but rather, other concurrent mental processes for task execution (although they are unlikely contingent negative variations; see *SI Discussion*). DC potentials likely reflect higher states of arousal, alertness, and or attention (Kovac et al. 2018). Indeed, low-frequency (delta band) EEG responses are closely linked to sustained attention (Kirmizi-Alsan et al. 2006). Interestingly, suppression of sustained responses during the delay period is reduced in individuals that rely on a covert rehearsal strategies (Linke et al. 2011). Thus, stronger AC responses for lower set sizes may reflect more covert rehearsal by our participants to refresh short-term memory representations from being overwritten and protecting memory items from decay. This interpretation is consistent with our brain-behavior correlations, which showed listeners with stronger reduction (cf. suppression) of sustained AC activity had higher *K* capacity limits (e.g., Fig. 3d). Larger neural responses for lower set sizes could reflect a stronger deployment of attention in easier, less taxing conditions.

Seminal studies in primates and fMRI in humans have demonstrated that neurons maintain a representation of the stimulus via sustained activity in prefrontal cortex which outlasts the eliciting event (Courtney et al. 1997). Sustained maintenance activation is observed across multiple brain regions in tasks involving various sensory modalities (Bashivan et al. 2014b; Yue et al. 2019). In agreement with prior neuroimaging WM studies (Grimault et al. 2014; Huang et al. 2016; Vogel and Machizawa 2004), we similarly found sustained EEG activity during WM maintenance is modulated by stimulus load. Our results corroborate fMRI findings (Kumar et al. 2016) by confirming successful auditory WM depends on elevated activity within AC. These structures are typically associated with sensory-perceptual processing rather than higher cognition (Boran et al. 2019; Yue et al. 2019). Thus, our data support an “embedded” rather than “buffer” account of WM, whereby auditory cortex acts to both encode perceptually relevant sound features but also functions as memory buffer to maintain information over short periods (cf. Yue et al. 2019).

We also found a stark hemispheric asymmetry in how AC predicts WM capacity. “High modulators” of left hemisphere AC activity showed superior auditory WM performance. In contrast, right hemisphere did not predict behavior. fMRI studies similarly show left AC activation positively correlates with behavioral performance in WM tasks (Kumar et al. 2016). Thus, while activity is bilateral during WM (current study; Grimault et al. 2014; Huang et al. 2016; Kumar et al. 2016), left hemisphere best correlates with behavioral capacity. Leftward laterality is perhaps expected given the well-known dominance of left cerebral hemisphere to auditory-linguistic processing (Hickok and Poeppel 2007). Indeed, both children and adults show strong brain asymmetry in WM organization, with greater rightward bias for spatial WM and leftward bias for verbal WM (Thomason et al. 2009). Given our WM task required covert verbal labeling, the stronger activation and association we find between left AC and behavior is consistent with the left hemisphere dominance of auditory-linguistic processing (Hickok and Poeppel 2007).

## Conflict of interest

none declared

## Acknowledgements

Work supported by the National Institute on Deafness and Other Communication Disorders of the National Institutes of Health (R01DC016267).

## Notes

### Competing Interest Statement

The authors have declared no competing interest.

